# Septin 2/6/7 complexes tune microtubule plus end growth and EB1 binding in a concentration- and filament-dependent manner

**DOI:** 10.1101/638296

**Authors:** Konstantinos Nakos, Megan R. Radler, Elias T. Spiliotis

## Abstract

Septins are filamentous GTP-binding proteins, which affect microtubule (MT) dependent functions including membrane trafficking and cell division, but their precise role in MT dynamics is poorly understood. Here, in vitro reconstitution of MT dynamics with SEPT2/6/7, the minimal subunits of septin heteromers, shows that SEPT2/6/7 has a biphasic concentration-dependent effect on MT growth. Lower concentrations of SEPT2/6/7 enhance MT plus end growth and elongation, while higher and intermediate concentrations inhibit and pause plus end growth, respectively. We show that SEPT2/6/7 has a 1.5-fold preference for GTP-over GDP-bound MT lattice, and competes with EB1 for binding to GTPγS-stabilized MTs, which mimic the EB1-preferred GDP-Pi state of polymerized tubulin. Strikingly, SEPT2/6/7 triggers EB1 dissociation from plus end tips in cis by binding to the MT lattice and in trans when MT plus ends collide with SEPT2/6/7 filaments. At these intersections, SEPT2/6/7 filaments were more potent barriers than actin filaments in pausing MT growth and dissociating EB1 in vitro and in live cells. These data demonstrate that SEPT2/6/7 complexes and filaments can directly impact MT plus end growth and the tracking of plus end-binding proteins, and thereby may facilitate the capture of MT plus ends at intracellular sites of septin enrichment.

**Highlight Summary for eTOC:** Knowledge of septin roles in MT dynamics is poor and confounded by knockdown studies. Here, in vitro reconstitution assays show concentration-dependent effects of SEPT2/6/7 on MT plus end growth, pausing and EB1 tracking. We found that SEPT2/6/7 filaments are potent than actin in pausing MT growth and dissociating EB1 from intersecting plus ends.

## Introduction

Septins are GTP-binding proteins that assemble into filamentous higher-order oligomers and polymers comprising a major component of the mammalian cytoskeleton along actin, microtubules and intermediate filaments (Spiliotis, 2018). Septins associate with microtubules (MTs) and affect MT organization and dynamics in various cell types (Kremer *et al.*, 2005; Bowen *et al.*, 2011; Ageta-Ishihara *et al.*, 2013; Bai *et al.*, 2013). Notably, septins are functionally involved in MT-dependent functions such as cell division and polarized membrane traffic (Spiliotis *et al.*, 2005; Spiliotis *et al.*, 2008; Estey *et al.*, 2010; Menon *et al.*, 2014; Karasmanis *et al.*, 2018). A growing number of studies suggests that septins are involved in the entry and growth of MTs in membrane protrusions (e.g., filopodia, cilia, micro-tentacles) (Ghossoub *et al.*, 2013; Nolke *et al.*, 2016; Ostevold *et al.*, 2017), but how septins affect the dynamic instability of MTs is not well understood.

Mammalian septins are a family of 13 paralogous genes, which are classified in four groups named after SEPT2, SEPT3, SEPT6 and SEPT7 (Kinoshita, 2003b; Weirich *et al.*, 2008; Russell and Hall, 2011). The minimal building block of septin heteromers is a palindromic hetero-hexamer that contains septins from the SEPT2, SEPT6 and SEPT7 groups in a stoichiometry of 2:2:2, which can further expand into a hetero-octamer upon addition of a septin paralog from the SEPT3 group (Kinoshita, 2003a; Sheffield *et al.*, 2003; Sirajuddin *et al.*, 2007; Kim *et al.*, 2011; Sellin *et al.*, 2011). Depending on the relative expression of septin paralogs, which differs among cell types, subunits of septin heteromers have been reported to colocalize with MTs (Spiliotis *et al.*, 2008; Spiliotis, 2010; Sellin *et al.*, 2012, 2014; Spiliotis, 2018; Targa *et al.*, 2019). SEPT9 binds MTs directly through N-terminal repeat motifs (K/R-R/x-x-D/E), which associate with the C-terminal tails of β-tubulin (Bai *et al.*, 2013), but it is unclear whether these motifs function similarly for septins of the SEPT6 and SEPT7 groups, which also associate with MTs; SEPT2 does not appear to bind MTs directly (Bai *et al.*, 2013).

How septins, as individual subunits or higher order complexes, impact the dynamic properties of MTs has been difficult to determine owing to disparate results from septin knock-down approaches. SEPT7 was initially posited to enhance MT dynamics, because SEPT7 knock-down increased MT acetylation, a marker of MT stability (Kremer *et al.*, 2005; Janke and Montagnac, 2017). However, this effect was attributed to cytosolic SEPT7, which sequestered the microtubule-associated protein 4 (MAP4) away from MTs (Kremer *et al.*, 2005) and promoted the interaction of histone deacetylase HDAC6 with cytosolic free tubulin (Ageta-Ishihara *et al.*, 2013). Comprehensive analysis of MT dynamics in SEPT2-depleted MDCK cells revealed a distinct increase in MT catastrophe, indicating that septins promote persistent MT growth and thereby, may promote MT growth (Bowen *et al.*, 2011). More recently, septins were proposed to promote the formation of MT-driven membrane protrusions through an interplay with the MT plus end tracking protein EB1 in cells that are infected with a *C. difficile* toxin (Nolke *et al.*, 2016). Despite these findings, it remains unclear how precisely septins regulate MT dynamics. To further our understanding, we reconstituted MT plus end and EB1 dynamics in vitro with soluble SEPT2/6/7 complexes and SEPT2/6/7 filaments.

## Results

### SEPT2/6/7 tunes MT plus end dynamics in a concentration-dependent manner

Using a TIRF microscopy assay that visualizes the polymerization and depolymerization of Rhodamine-labeled tubulin onto stable HiLyte 488-labeled MT seeds, we tested how MT dynamics are impacted by different concentrations of recombinant SEPT2/6/7 complexes, which were purified from bacteria. Quantification of the parameters of MT dynamics revealed a biphasic concentration-dependent response for catastrophe frequencies and MT lengths (Figure 1, C and D). Lower concentrations of SEPT2/6/7 (10 - 400 nM) reduced catastrophe frequencies by 20- 34% and increased MT lengths by 1.2-to-1.8-fold, whereas higher concentrations of SEPT2/6/7 (>1 μM) had the opposite effect, enhancing catastrophe frequencies by 1.2-to-1.6-fold and decreasing MT lengths by 30-48%. A similar, but less pronounced, biphasic effect was observed for the rates of MT growth, which increased slightly at 10 nM of SEPT2/6/7 and decreased gradually at concentrations >800 nM, while no effect was observed between 10 and 800 nM (Figure 1A). Interestingly, the rates of MT depolymerization were not affected in a biphasic manner, but rather decreased steadily with increasing concentrations of SEPT2/6/7 (Figure 1B). As also evidenced by the kymographs of MT dynamics (Figure 1, E-M), these data show that lower concentrations of SEPT2/6/7 (<0.5 μM) induce persistent MT growth by decreasing the frequency of catastrophe events and rates of depolymerization. However, micromolar concentrations of SEPT2/6/7 (>1 μM) dampen MT dynamics and reduce MT lengths by decreasing rates of MT growth and enhancing catastrophe.

**Figure 1.**
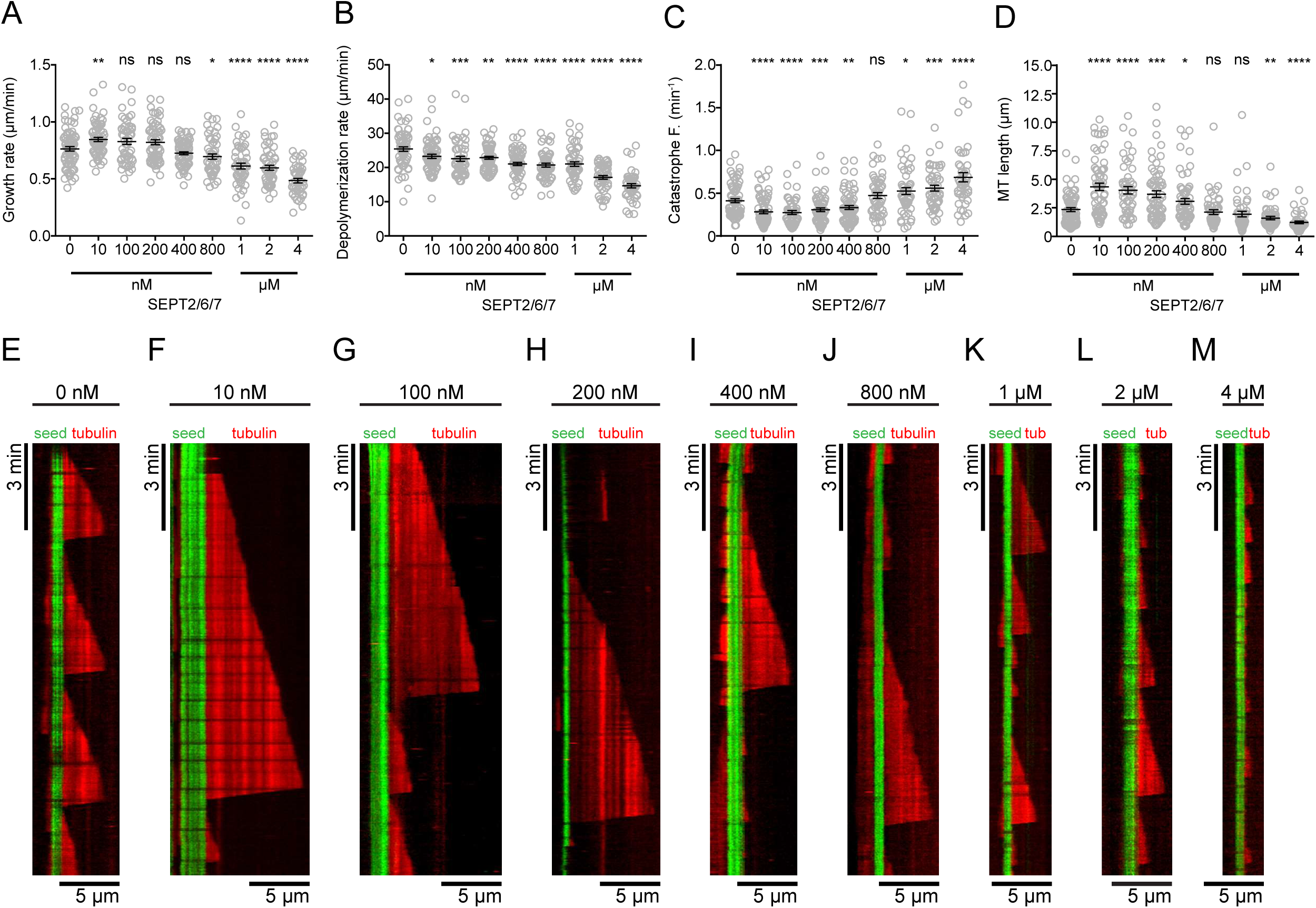
SEPT2/6/7 tunes MT dynamics in a concentration-dependent manner. (A-D) Dot plots show mean (± SEM) rates of MT plus end polymerization (A) and depolymerization (B), and frequency of MT plus end catastrophe (C) and length (D) with SEPT2/6/7 at concentrations of 0 nM (n = 69), 10 nM (n = 65), 100 nM (n = 51), 200 nM (n = 61), 400 nM (n = 66), 800 nM (n = 50), 1 µM (n = 51), 2 µM (n = 49) and 4 µM (n = 44). (E-M) Kymographs show representative microtubule plus end dynamics (red) upon nucleation from MT seeds (green) in the presence of 0 nM (E), 10 nM (F), 100 nM (G), 200 nM (H), 400 nM (I), 800 nM (J), 1 µM (K), 2 µM (L) and 4 µM (M) of SEPT2/6/7. ns: non-significant (*p* > 0.05). *p < 0.05, **p < 0.01, ***p < 0.001, ****p < 0.0001.

Because SEPT2/6/7 complexes assemble into higher-order filamentous structures, which depend on concentration and ionic conditions, we reasoned that the biphasic effect might be due to concentration-dependent differences in the assembly state of SEPT2/6/7 complexes. To assess the higher-order state of SEPT2/6/7, we performed a sedimentation assay after diluting and incubating SEPT2/6/7 complexes in the buffer of the MT dynamics assay. Quantifications of SEPT2/6/7 in the pellet and supernatant fractions showed that the levels of sedimenting higher-order septins were fairly equivalent up to concentrations of 400 nM, but increased in a stepwise concentration-dependent manner above 800 nM (Figure S1, A and B). This increase in the amount of higher-order SEPT2/6/7 correlated with the dampening of MT dynamics observed at concentrations >800 nM (Figure 1, A and C-D). Hence, higher order fibrillar SEPT2/6/7 does not appear to favor MT growth and elongation, which is enhanced by lower concentrations of SEPT2/6/7 (Figure 1).

Given that MT growth and dynamics are inhibited by higher concentrations of SEPT2/6/7 (>800 nM), we reasoned that higher order SEPT2/6/7 may associate with free unpolymerized tubulin, lowering the effective concentration of tubulin that is available to polymerize. This is plausible as we recently found that SEPT9 binds and recruits free tubulin to the MT lattice (Nakos *et al.*, 2019). To test this possibility, SEPT2/6/7 and SEPT9 complexes were immobilized on functionalized glass chambers and fluorescent tubulin was added in the buffer conditions of the MT dynamics assay (Figure S2A). Binding of soluble tubulin to septin complexes was visualized by TIRF imaging similar to a previous assay of MT nucleation (Lazarus *et al.*, 2013). While tubulin was increasingly bound to chambers with sub-micromolar concentrations of SEPT9, no binding was observed for a similar concentration range of SEPT2/6/7 (Figure S2, B and C). These data indicate that SEPT2/6/7 does not sequester unpolymerized tubulin and the effects of SEPT2/6/7 on MT dynamics depend on the septin concentration and assembly state (oligomers vs. higher order polymers).

### SEPT2/6/7 pauses MT plus end growth

In kymographs of MT dynamics, we noticed that MT plus ends exhibited a variety of pausing events in the presence of SEPT2/6/7 complexes. In several kymographs, MT plus ends paused transiently while growing and subsequently, depolymerized or continued to grow before undergoing catastrophe (Figure 2, A-B and D; arrowheads). Although much less frequent, MT plus ends also paused transiently during catastrophe before depolymerizing to the MT seed (Figure 2, C-D and G). Additionally, the slope of depolymerization events was shallower, which was consistent with decreased rates of depolymerization (Figure 1).

**Figure 2.**
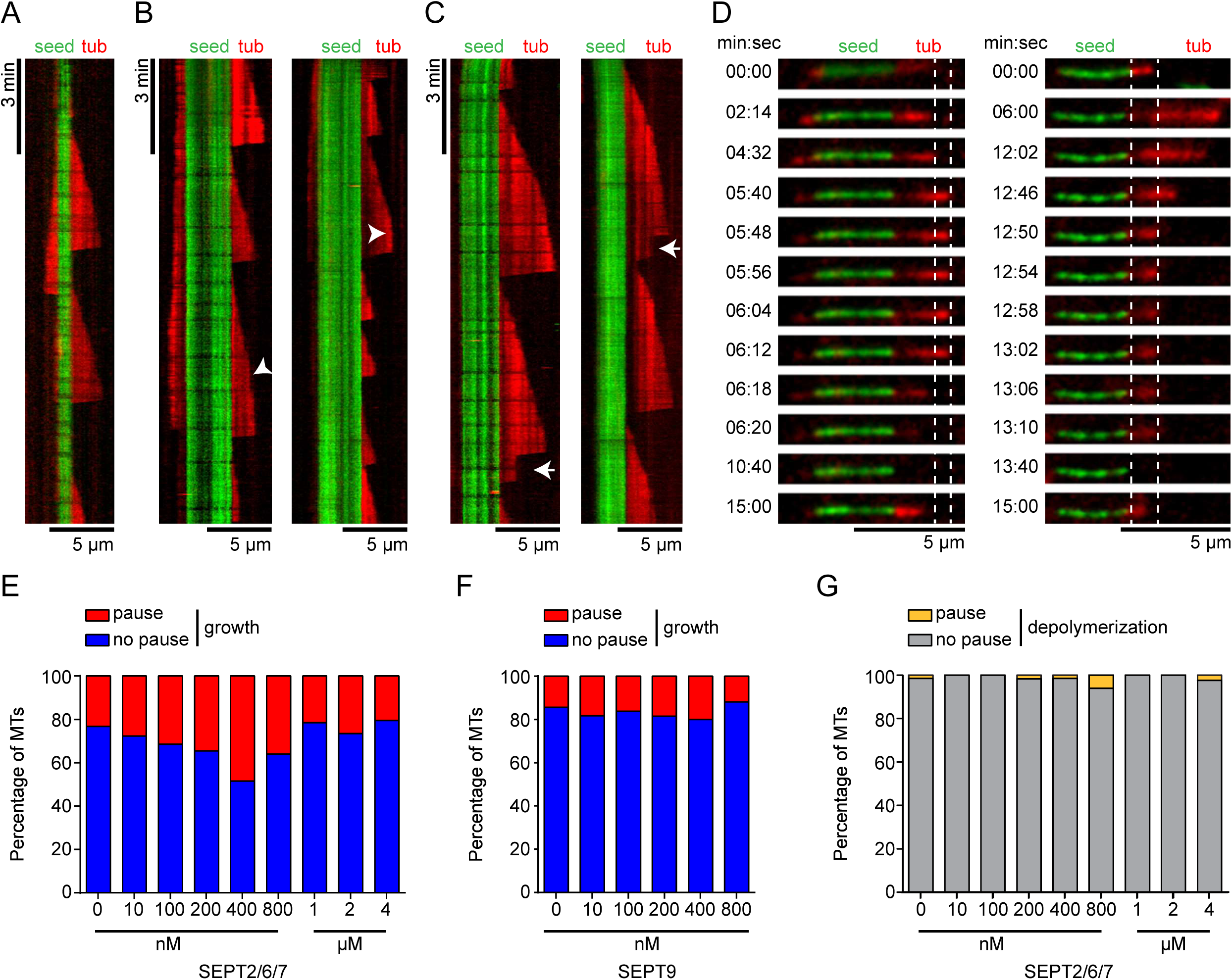
SEPT2/6/7 pauses MT plus end growth. (A-C) Kymographs show pausing events during MT growth (B) or shrinkage (C) in the presence of 0 nM (A), 400 nM (B and C, left) and 800 nM (B and C, right) of SEPT2/6/7. (B) Arrowheads point to pause events in the presence of 400 nM and 800 nM of SEPT2/6/7. (C) Arrows point to pause events observed during depolymerization of MT plus ends in the presence of 400 nM and 800 nM of SEPT2/6/7. (D) Images show still frames of MT seeds (green) and dynamic MTs (red) undergoing a pause event during growth in the presence of 800 nM SEPT2/6/7 (D, left) or a pause event during depolymerization in the presence of 400 nM SEPT2/6/7 (D, right). Dashed lines indicate MT length during pause events. Time is in min:sec. (E-F) Bar graphs show the percentage of MTs with continuous growth (no pause, blue) or pause while growing (pause, red) in the presence of 0 nM (n = 69), 10 nM (n = 65), 100 nM (n = 51), 200 nM (n = 61), 400 nM (n = 66), 800 nM (n = 50), 1 µM (n = 51), 2 µM (n = 49) and 4 µM (n = 44) of SEPT2/6/7, or in the presence of 0 nM (n = 21), 10 nM (n = 44), 100 nM (n = 31), 200 nM (n = 38), 400 nM (n = 40) and 800 nM (n = 34) of SEPT9_i1. (G) Bar graph shows the percentage of MTs with continuous depolymerization (no pause, grey) or pause during shrinkage (pause, orange) in the presence of 0 nM (n = 69), 10 nM (n = 65), 100 nM (n = 51), 200 nM (n = 61), 400 nM (n = 66), 800 nM (n = 50), 1 µM (n = 51), 2 µM (n = 49) and 4 µM (n = 44) of SEPT2/6/7.

Given the concentration dependence of SEPT2/6/7 effects on MT dynamics, we analyzed the frequency of pausing events that occurred in growth phases. Strikingly, MT pausing increased with increasing concentrations of SEPT2/6/7 peaking at 400 nM, where the percentage of MTs with pausing events doubled from 23% to 48% (Figure 2E). This pausing effect was unique to SEPT2/6/7 as SEPT9 did not cause a similar effect (Figure 2F). However, at SEPT2/6/7 concentrations higher than 400 nM, MT pausing began to decrease and micromolar concentrations of SEPT2/6/7 did not increase pausing above the levels observed in the absence of SEPT2/6/7. Hence, MT pausing appears to be an intermediate phenotype that occurs by SEPT2/6/7 complexes in between concentrations that promote and inhibit MT growth and elongation.

### SEPT2/6/7 associates with the MT lattice and plus ends

Based on the effects of SEPT2/6/7 on MT plus end growth and pausing, we sought to examine whether SEPT2/6/7 complexes associate with the lattice and/or tips of dynamic MT plus ends. At low concentrations (10-100 nM), mCherry-SEPT2/6/7 localized sparsely and in patches on the lattice of MT seeds and growing plus ends, and occasionally at the tips of plus ends (Figure 3A). In kymographs of time-lapse imaging, we observed binding of mCherry-SEPT2/6/7 to the lattice of the MT plus end segments, which persisted for several seconds to a few minutes, and to the tips of polymerizing and depolymerizing plus ends (Figure 3A). Because of the sparse localization, we quantified MTs categorically based on the distribution of mCherry-SEPT2/6/7 (Figure 3B). At 10 nM, mCherry-SEPT2/6/7 localized to the lattice of the seed and/or plus end segments of most MTs (~80%), and the tips of plus ends on 27% of total MTs. At 100 nM, however, binding of mCherry-SEPT2/6/7 to plus end tips was rare, and SEPT2/6/7 associated exclusively with the lattice of MT seeds (65%) or with both the lattice of MT seeds and plus end segments (Figure 3B). At concentrations >100 nM, mCherry-SEPT2/6/7 decorated MTs more extensively (Figure 3C). Quantification of mCherry-SEPT2/6/7 fluorescence on the lattice of MT seeds versus plus ends showed a 1.5-fold preference for the GMPCPP-stabilized lattice (Figure 3, C and D), which is significantly less than the ~4-fold difference that was previously observed for SEPT9, which in similar concentrations (400-800 nM) did not fully decorate the lattice segments of MT plus ends (Nakos *et al.*, 2019). Collectively, our results show that SEPT2/6/7 has a higher affinity than SEPT9 for the GDP-bound lattice of MT plus end segments.

**Figure 3.**
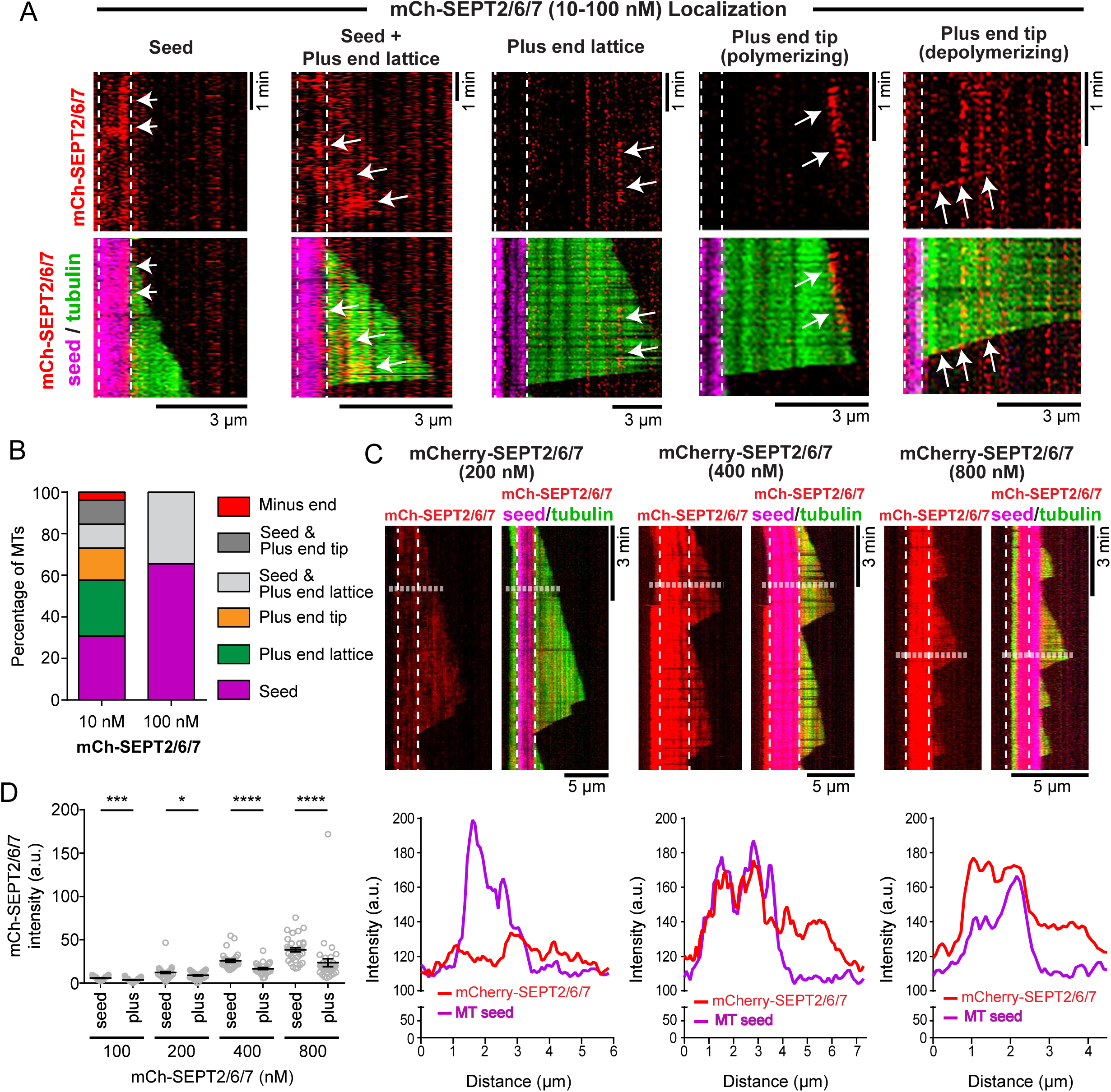
SEPT2/6/7 associates with MT lattices and plus ends. (A) Kymographs show examples of mCherry-SEPT2/6/7 (red; arrows) localization to the MT lattice of GMPCPP seeds (magenta; vertical dashed lines) and plus end segments (green) as well as to the tips of polymerizing and depolymerizing plus ends. (B) Bar graph shows the percentage of MTs, which contained mCherry-SEPT2/6/7 (n = 26-29) only on the lattice of GMPCPP-stabilized MT seeds (magenta) and plus end segments (green) or only on plus end tips (orange). In addition, percentage of MTs with mCherry-SEPT2/6/7 on minus ends (red) and on both seeds and plus end lattice (light gray) or tips (dark grey) were quantified. (C) Kymographs show the localization of mCherry-SEPT2/6/7 (red) at 200 nM, 400 nM and 800 nM. Note that mCherry-SEPT2/6/7 decorates the MT lattice of stable GMPCPP MT seeds (magenta or dashed outlines) and dynamic plus end segments (green). Line scans show the fluorescence intensity of mCherry-SEPT2/6/7 (red line) and GMPCPP-stabilized MT seed (HiLyte-647-tubulin; magenta line) along MT segments, which are outlined in kymographs with horizontal dashed lines. (D) Dot plots show the mean (± SEM) fluorescence intensity of mCherry-SEPT2/6/7 on GMPCCP-stable seeds and the lattice as well as tips of dynamic plus end segments. Quantification was performed from images of MTs after 15 minutes of incubation with 100 nM (n = 29), 200 nM (n = 35), 400 nM (n = 30) and 800 nM (n = 35) of mCherry-SEPT2/6/7. ns: non-significant (*p* > 0.05). *p < 0.05, **p < 0.01, ***p < 0.001, ****p < 0.0001.

### SEPT2/6/7 complexes inhibit MT plus end binding and tracking of EB1

Recent reports of an interaction between EB1 and septins (Nolke *et al.*, 2016) raises the question of whether EB1 and SEPT2/6/7 cooperate or compete with one another on MT plus ends. Given the effects of SEPT2/6/7 on MT plus ends, we sought to test whether SEPT2/6/7 affects the binding and dynamics of the plus end-binding protein EB1, which can also provide more insight into how SEPT2/6/7 impacts MT plus end dynamics.

We reconstituted MT dynamics in the presence of recombinant EB1-GFP and mCherry-SEPT2/6/7 (Figure 4A). Tracking of EB1-GFP on MT plus ends required buffer conditions (50 mM KCl), which limit diffusive interactions of EB1-GFP with the MT lattice (Zanic *et al.*, 2009). In these ionic conditions, the MT-binding affinity of mCherry-SEPT2/6/7 was reduced, but mCherry-SEPT2/6/7 exhibited the same concentration-dependent pattern of decoration with a modest preference for GMPCPP-bound MT seeds (Figure 4A).

**Figure 4.**
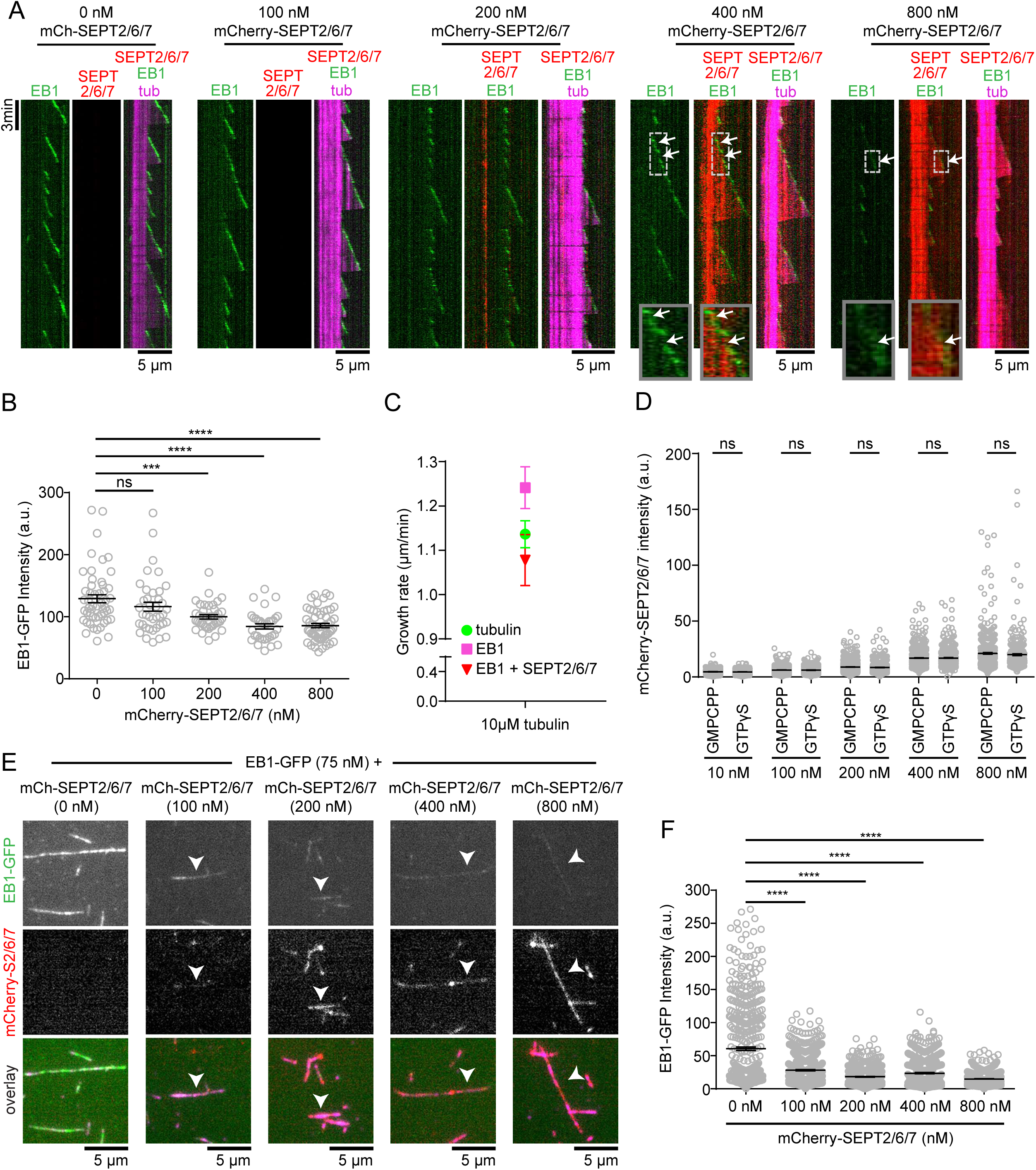
SEPT2/6/7 inhibits EB1 binding to MT plus ends in a concentration-dependent manner.(A) Kymographs show examples of polymerizing MTs (magenta; GMPCCP-stable seeds in bright magenta) with EB1-GFP (75 nM; green) in the absence or presence of increasing concentrations of mCherry-SEPT2/6/7 (red; 100 - 800 nM). Note that the intensity of EB1-GFP decreases with increasing concentrations of mCherry-SEPT2/6/7. Insets show in higher magnification the binding of mCherry-SEPT2/6/7 to plus end sites proximal to EB1-GFP and the persistence of mCherry-SEPT2/6/7 to these sites (arrows) while plus end continue to grow with newly added EB1-GFP. The fluorescence intensity of mCherry-SEPT2/6/7 at 800 nM was adjusted as it is saturated when shown in the same dynamic range and exposure times with concentrations below 800 nM. (B) Mean (± SEM) fluorescence intensity of EB1-GFP at MT plus end tips in the absence (n = 52) or presence of 100 nM (n = 39), 200 nM (n = 39), 400 nM (n = 33) and 800 nM (n = 54) mCherry-SEPT2/6/7. (C) Graph shows mean (± SEM) growth rates of tubulin in the absence of mCherry-SEPT2/6/7 or EB1-GFP (green; n = 32), and in the presence of 75 nM GFP-EB1 (magenta; n = 28) without or with 800 nM mCherry-SETP2/6/7 (red; n = 21). Note that the enhancement of MT growth rate by EB1-GFP is inhibited by mCherry-SEPT2/6/7. (D) Dot plots show the mean (± SEM) fluorescent intensity of mCherry-SEPT2/6/7 on the lattice of GMPCPP- and GTPγS-stabilized MTs. Quantification was performed from static images of MTs after 15 min of incubation in the presence of 10 nM (GMPCPP, n = 587; GTPγS, n = 482), 100 nM (GMPCPP, n = 597; GTPγS, n = 511), 200 nM (GMPCPP, n = 730; GTPγS, n = 537), 400 nM (GMPCPP, n = 614; GTPγS, n = 457) and 800 nM (GMPCPP, n = 454; GTPγS, n = 322) of mCherry-SEPT2/6/7. (E) Images show examples of GTPγS seeds (magenta) incubated with 75 nM EB1-GFP (green) and increasing concentrations of mCherry-SEPT2/6/7 (red; 100 - 800 nM). Note that EB1-GFP binding decreases with increasing concentrations of mCherry-SEPT2/6/7. (F) Mean (± SEM) fluorescent intensity of EB1-GFP on GTPγS-stabilized MTs in the absence or presence of mCherry-SEPT2/6/7. Quantification was performed from static images of MTs after 15 minutes of incubation without (n = 528) or with mCherry-SEPT2/6/7 at concentrations of 100 nM (n = 626), 200 nM (n = 559), 400 nM (n = 581) and 800 nM (n = 626). ns: non-significant (*p* > 0.05). *p < 0.05, **p < 0.01, ***p < 0.001, ****p < 0.0001.

In the absence of mCherry-SEPT2/6/7, EB1-GFP (75 nM) tracked on growing MT plus ends and was absent from MT tips during catastrophe as previously reported (Bieling *et al.*, 2007; Bieling *et al.*, 2008; Dixit *et al.*, 2009). In kymographs, EB1-GFP appeared as diagonal lines, outlining the trajectory of the growing MT plus ends (Figure 4A). Strikingly, the fluorescence intensity of EB1-GFP decreased with increasing concentrations of mCherry-SEPT2/6/7, which was indicative of a reduction in the binding of EB1-GFP to growing MT plus ends (Figure 4A). Quantification of EB1-GFP fluorescence showed a statistically significant reduction from the plus ends of MTs in the presence of 400-800 nM mCherry-SEPT2/6/7 (Figure 4B). Moreover, SEPT2/6/7 had a dominant negative effect on the enhancement of plus end growth rates by EB1-GFP (Figure 4C); an increase in rates of plus end growth by EB1 is consistent with previous studies (Vitre *et al.*, 2008; Zanic *et al.*, 2009; Zanic *et al.*, 2013).

Inhibition of EB1-GFP binding and tracking by SEPT2/6/7 pointed to a few possibilities. First, EB1-GFP might be sequestered by soluble SEPT2/6/7 complexes as has been shown before for tau (Ramirez-Rios *et al.*, 2016). Second, SEPT2/6/7 may compete transiently with EB1 for binding to tips of MT plus ends that transition from tubulin-GTP to tubulin-GDP-Pi, which are respectively the low- and high-affinity binding sites of EB1 (Zanic *et al.*, 2009; Maurer *et al.*, 2011; Maurer *et al.*, 2012). Third, binding of SEPT2/6/7 to the GDP-bound MT lattice may cause long-range conformational changes along MT protofilaments, affecting allosterically the binding of EB1 to plus ends (Zanic *et al.*, 2013; Brouhard and Rice, 2018).

To examine whether EB1-MT binding is inhibited through a direct interaction between SEPT2/6/7 and EB1, we performed protein-binding assays with EB1 and SEPT2/6/7 in buffers of two different ionic conditions including those of the TIRF in vitro assay. In neither of these conditions, SEPT2/6/7 interacted with EB1 (Figure S3A). In contrast, we were able to detect an association between EB1 and SEPT5 (Figure S3, B and C), a septin that possess the EB1-interaction motif SxIP (Honnappa *et al.*, 2009; Jiang *et al.*, 2012). Hence, SEPT2/6/7 is unlikely to inhibit EB1-MT binding through a direct interaction.

Next, we tested the possibility of a competition between EB1 and SEPT2/6/7 for binding to MTs using GMPCPP- and GTPγS-stabilized MTs, which respectively mimic the GTP-bound (low EB1 affinity) and GDP-Pi-bound (high EB1 affinity) conformations of polymerized tubulin (Zanic *et al.*, 2009; Maurer *et al.*, 2011; Maurer *et al.*, 2012; Roth *et al.*, 2018). Unlike EB1, SEPT2/6/7 showed no preference between GMPCPP- and GTPγS-stabilized MTs (Figure 4D), indicating that SEP2/6/7 interacts with GTPγS-bound MT lattices as optimally as observed with the GMPCPP-stabilized MT seeds (Figure 3, B-D). In a competition experiment, EB1-GFP (75 nM) was incubated together with increasing concentrations of mCherry-SEPT2/6/7 in the presence of GTPγS-stabilized MTs. Decoration of MTs with EB1-GFP was reduced and SEPT2/6/7 accumulated to MTs with concomitant displacement of EB1-GFP (Figure 4, E and F). These data suggest that association of SEPT2/6/7 with the GDP-Pi lattice of MT plus ends could prevent binding and tracking of EB1.

Despite the competition between EB1 and SEPT2/6/7 for binding to the lattice of GTPγS-stabilized MTs, which mimic the GDP-Pi segments of MT plus end tips (high affinity state of EB1) (Maurer *et al.*, 2011), we were not able to resolve any exchange between mCherry-SEPT2/6/7 and EB1-GFP on the tips of growing plus ends – mCherry-SEPT2/6/7 and EB1-GFP did not alternate tracking on plus end tips (Figure 4A). Occasionally, mCherry-SEPT2/6/7 localized near the tips of plus ends transiently or remained at the same position while the plus ends continued to grow with EB1-GFP (arrows, insets in Figure 4A). This might be due to the higher affinity of EB1 for GDP-Pi than SEPT2/6/7; note that full decoration of GTPγS MTs by EB1 is achieved in much lower concentrations than SEPT2/6/7 (Figure 4E). However, even at high concentrations (800 nM) of SEPT2/6/7, which strongly diminish EB1 binding to plus ends, we did not observe a distinct accumulation of SEPT2/6/7 to plus end tips – SEPT2/6/7 presence near the plus end tips was indistinguishable from its localization along the MT lattice (Figure 4A). Thus, while a direct competition at the plus end tips cannot be ruled out, we posit that MT-bound SEPT2/6/7 may also affect EB1 binding by causing long-range conformational changes on MT protofilaments (Brouhard and Rice, 2018).

### End-on collisions of MT plus ends with SEPT2/6/7 filaments results in pausing and EB1 dissociation

Although SEPT2/6/7 does not track with growing MT plus ends in cells or in vitro, MT plus ends have been observed to collide end-on with septins (Bowen *et al.*, 2011; Nolke *et al.*, 2016). We, therefore, sought to determine whether septin filaments can directly impact MT plus end dynamics during end-on encounters, which do not involve association of septin complexes with the MT lattice.

We reconstituted MT dynamics in the presence of septin filaments by immobilizing mCherry-SEPT2/6/7 and MT seeds on glass substrata. Upon introduction of soluble tubulin, we were able to visualize the polymerizing plus ends of MTs intersecting with filamentous septins. During these encounters, MT plus ends paused, continued to grow or depolymerized (Figure 5A). Quantification showed that ~46% of encounters with immobilized mCherry-SEPT2/6/7 filaments resulted in MT plus pausing, while continuation of growth and depolymerization were observed at ~43% and ~9% of intersections, respectively. Notably, MT pausing events were significantly higher (~2-fold) compared to plus end collisions with immobilized actin filaments (Figure 5B). Upon encountering F-actin, continuation of MT plus end growth was much more frequent than pausing – continued growth (73%) was 3-fold more frequent than pausing (~23%) (Figure 5B).

**Figure 5.**
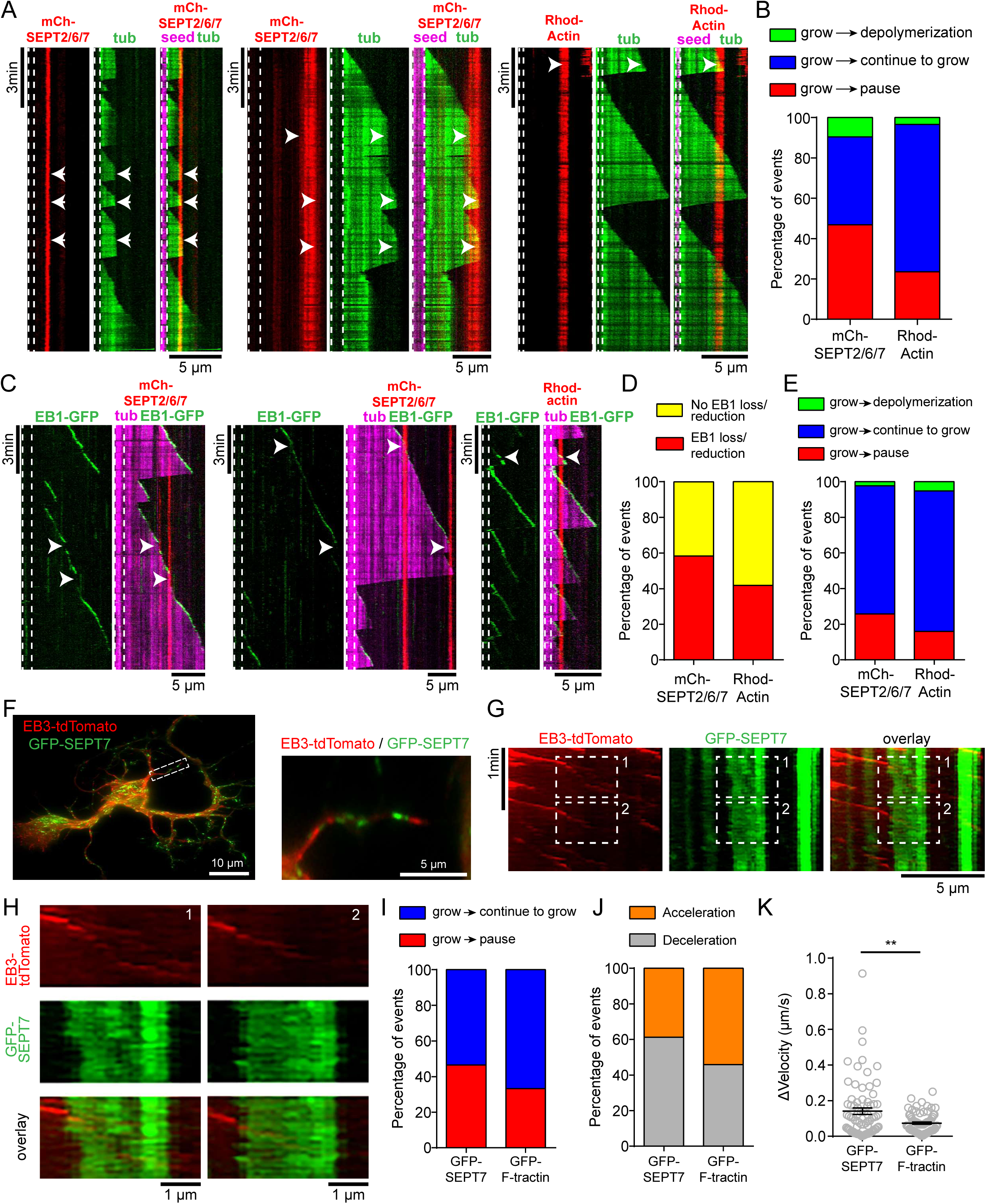
MT plus end collisions with SEPT2/6/7 filaments enhance pausing and EB1 dissociation. (A) Kymographs show examples of end-on collisions of growing MT plus ends (green; GMPCPP-stable MT seeds in magenta) with mCherry-SEPT2/6/7 or Rhodamine-labeled actin filaments (red), which were immobilized on the glass matrix. Arrowheads point to pauses of MT plus ends at intersections with mCherry-SEPT2/6/7 or Rhodamine-labeled actin filaments. (B) Bar graph shows the percentage of intersection events in which MT plus ends paused (red) upon encountering SEPT2/6/7 (n = 207 events) or actin filaments (n = 178 events), continued to grow (blue) or depolymerized (green). Note that the percentage of pause events increased by two-fold upon encountering SEPT2/6/7 compared to actin. (C) Kymographs show examples of dynamic MTs (magenta) assembled with 75 nM EB1-GFP (green) in the presence of immobilized mCherry-SEPT2/6/7 or Rhodamine-labeled actin filaments (red). Arrowheads point to loss or diminution of EB1-GFP from plus end tips at intersections with septin and actin filaments. Note that EB1-GFP binding at the MT plus end tips is reduced or lost at points of intersection with mCherry-SEPT2/6/7 filaments. (D) Percentage of collision events during which EB1-GFP was reduced or lost from MT plus ends (red) or remained unaffected (yellow) upon encountering mCherry-SEPT2/6/7 (n = 132 events) or actin filaments (n = 213 events). (E) Bar graph shows the percentage of events of MT plus ends with EB1-GFP (75 nM), which paused (red) upon encountering mCherry-SEPT2/6/7 (n = 132 events) or actin filaments (n = 213 events), continued to grow (blue) or depolymerized (green). Note that the percentage of pause events is higher upon encountering SEPT2/6/7 (26%) than actin filaments (16%). (F-H) Still images (F) show a rat hippocampal neuron (DIV3) co-transfected with plasmids expressing GFP-SEPT7 and EB3-tdTomato. An outlined neurite region (dashed rectangle) in which EB3-tdTomato movement was analyzed with respect to GFP-SEPT7 is shown in higher magnification and corresponds to the depicted kymographs (G-H). Kymographs show the trajectories of EB3-tdTomato comets (red) that intersect with GFP-SEPT7 (green) (G). Outlined regions with diminution of EB3-tdTomato intensity (dashed rectangles 1 and 2) during overlap with GFP-SEPT7 and thereafter resumed movement (dashed rectangle 2) are shown in higher magnification (H). (I) Bar graph shows the percentage of EB3 comets which paused (red) or continued to move (blue) upon encountering SEPT7 (n = 342 comets; 27 neurites) or actin filaments (n = 399 comets; 22 neurites). Note that the percentage of pause events is higher upon encountering SEPT7 (46%) than actin filaments (33%). (J) Bar graph shows the percentage of EB3 comets that resumed movement with reduced (deceleration; grey) or enhanced velocity (acceleration; orange) compared to their velocity prior to pausing at SEPT7 (n = 124) or actin filaments (n = 168). Deceleration was more frequent after pausing at SEPT7 (61%) than actin filaments (46%). (K) Mean (± SEM) change of velocity in the events of deceleration of EB3 comets after pausing at SEPT7 (n = 76) or actin filaments (n = 77). ns: non-significant (*p* > 0.05). *p < 0.05, **p < 0.01, ***p < 0.001, ****p < 0.0001.

Next, we asked whether MT plus ends exhibited similar behavior upon encountering septin and actin filaments in the presence of EB1-GFP. Interestingly, EB1-bound MT plus ends paused less than free MT plus ends at both septin and F-actin intersections (Figure 5, C and E). However, EB1-bound plus ends still paused more frequently at intersections with septins than actin filaments (26% vs. 16%) and dissociation of EB1 from plus ends was more frequent at collisions with SEPT2/6/7 than actin filaments (Figure 5, D and E). These data show that pausing of MT plus ends and EB1 dissociation is more likely to occur at intersections with septin than actin filaments.

To test whether septin filaments are a more potent barrier to MT plus end growth than actin filaments in living cells, we imaged and quantified the dynamics of EB3-tdTomato at sites of intersection with GFP-SEPT7 or GFP-F-tractin in primary rat hippocampal neurons (Figure 5, F-H). In agreement with the *in vitro* data, EB3 comets paused more frequently upon intersecting with GFP-SEPT7 than GFP-F-tractin (~46% vs ~33%; Figure 5I). Kymographs showed that EB3 comets became dimmer or disappeared upon contact with SEPT7 filaments (Figure 5, G and H). EB3 comets were also found to resume growth with reduced (deceleration) or enhanced (acceleration) rates compared to their velocity prior to pausing at SEPT7 or F-tractin. Notably, deceleration of MT plus end growth was more frequent after collision with SEPT7 than F-tractin (Figure 5J) and the reduction in velocity was two-fold higher upon encountering septin (ΔV = 0.14 ± 0.02 μm/s) than actin (ΔV = 0.07 ± 0.006 μm/s) (Figure 5K); frequency and rates of accelerated EB3 movement were not different between septin and actin. Thus, septin filaments not only are more potent barriers than actin filaments in pausing MT plus ends, but they also slow down MT re-growth after pausing.

## Discussion

A growing number of studies implicate septins in MT organization and dynamics as well as MT-dependent functions, but how septins impact the dynamic instability of MTs is not well understood. Results from cell biological experiments have been difficult to interpret as septins are multifunctional proteins with various binding partners, several of which associate with the cytoskeleton. Hence, in vitro reconstitution experiments can provide valuable insights by elucidating the direct effects of septins on MT dynamics.

In this study, MT dynamics were reconstituted with SEPT2/6/7 complexes and filaments. Our results show that the effects of SEPT2/6/7 on MT dynamics are dependent on their concentration and state of assembly, which may account for the disparity of previous findings obtained from septin knock-down approaches. We found a biphasic graded effect on MT dynamics which correlated with a concentration-dependent assembly of SEPT2/6/7 into higher-order filaments. At lower concentrations (10-100 nM), where assembly of SEPT2/6/7 complexes into higher-order filaments is limited, MT growth and elongation were enhanced. However, at higher concentrations (low μM), which enhance the filamentous state of SEPT2/6/7, MT growth was stunted. Interestingly, in the intermediate range of concentrations, SEPT2/6/7 functions as a pausing factor, causing transient arrests of MT plus end growth.

The effects of SEPT2/6/7 on MT dynamics have striking similarities and differences with SEPT9. While both SEPT2/6/7 heteromers and SEPT9 homomers promote the growing phase of MTs, SEPT2/6/7 achieves this effect at significantly lower concentrations than SEPT9 (Nakos *et al.*, 2019). Moreover, micromolar concentrations of SEPT9 stabilize elongated MTs, which seize to grow and shrink, while micromolar SEPT2/6/7 dampens MT dynamics without stabilizing MTs, which undergo more catastrophe. Additionally, SEPT9 binds and recruits unpolymerized tubulin to the lattice of MTs (Nakos *et al.*, 2019), while SEPT2/6/7 does not bind unpolymerized tubulin. Conversely, submicromolar concentrations of SEPT2/6/7 can pause MT plus end growth, which was not observed with SEPT9. Taken together, these data indicate that there are septin subunit-and paralog-specific differences in the modulation of MT dynamics, which might be physiologically relevant for the modulation of MT dynamics in cell types and pathological conditions with distinct patterns of septin expression.

Our results show that SEPT2/6/7 can trigger dissociation of EB1 from MT plus ends in *cis* by binding to the lattice of plus ends or *in trans* by coming in contact as a filamentous structure with on-coming MT plus ends. A similar effect has been shown for tau, which antagonizes EB1-binding to plus ends (Ramirez-Rios *et al.*, 2016). However, unlike tau which inhibits EB1-MT binding by forming a soluble complex with EB1 off MTs, SEPT2/6/7 does not appear to sequester EB1 off MTs. Interestingly, SEPT2/6/7 competes with EB1 for binding to GTPγS-stabilized MTs, which resemble the EB1-prefered tubulin-GDP-Pi conformation of plus end tips. In addition, SEPT2/6/7 is able to reduce the enhanced MT growth rates that we observed in the presence of EB1. Although EB1 dissociation upon MT plus end contact with SEPT2/6/7 filaments indicates a direct competition between EB1 and septins at tips of MT plus ends, SEPT2/6/7 tracking on MT plus ends is rare. Moreover, we have not been able to observe an exchange of EB1 and SEPT2/6/7 on MT plus ends. If SEPT2/6/7 does not affect EB1 binding at the tips of plus ends, binding of SEPT2/6/7 to the MT-GDP lattice may result in conformational changes, which are propagated to the tips of plus ends allosterically along the protofilaments of MTs (Brouhard and Rice, 2018). Evidence for this mechanism comes from studies of the potent MT polymerase XMAP215 and EB1 (Zanic *et al.*, 2013). Even though XMAP215 and EB1 do not interact, a synergistic effect is posited to occur through conformational changes (Zanic *et al.*, 2013; Brouhard and Rice, 2018). Functionally, a septin-mediated dissociation of EB1 from MT plus ends could provide a mechanism for pausing and stable capture of MT plus ends. Given that EB1 appears to protect MT plus ends from pausing during end-on collisions with septin and actin filaments (Figure 5), dissociation of EB1 would further enhance pausing of MT plus ends upon contact with septin filaments.

How are the concentration-dependent effects that septins exert on MT plus ends in vitro relevant to the concentration, organization and localization of septins in living cells? MT growth and elongation are enhanced at septin concentrations of hundreds of nanomolar, which reportedly is the estimated concentration of septins such as SEPT2 (~800 nM), SEPT6 (~200 nM), SEPT7 (~500 nM) and SEPT9 (~400 nM) in HeLa cells (Hein *et al.*, 2015). By contrast, micromolar septin concentrations dampen MT dynamics, resulting in stunted growth or complete MT stabilization. Given that septins are not uniformly distributed in cells and are enriched in specific subcellular regions (e.g., membrane domains of micron-scale curvature), it is plausible that septins tune MT dynamics in a region-specific manner according to their local concentration. Therefore, persistent MT growth could occur in cytoplasmic areas of moderate septin concentration, while MT pausing and stabilization could occur at cytoplasmic and membrane regions of septin enrichment. The latter could be key for the capture of MT plus ends at the base of membrane protrusions, where septins accumulate at domains of micron-scale curvature (Zhang *et al.*, 1999; Steels *et al.*, 2007; Akil *et al.*, 2016; Bridges *et al.*, 2016; Dolat and Spiliotis, 2016; Cannon *et al.*, 2019), or the stabilization and dampening of MT plus end dynamics at the tips of primary cilia, in which septins are also highly enriched (Hu *et al.*, 2010; Ghossoub *et al.*, 2013; Fliegauf *et al.*, 2014).

Modulation of MT plus end growth at sites of contact with septin filaments suggests that septins may function as cytoplasmic barriers, which gate spatially and temporally MT growth and invasion. Recent studies have shown that actin filaments act as physical barriers that regulate MT growth in vitro and in cultured cells (Colin *et al.*, 2018; Inoue *et al.*, 2019). Our results show that septins are more potent barriers in pausing and dampening MT re-growth than actin filaments. As septins associate preferentially with transverse arcs of migrating cells (Dolat *et al.*, 2014), they may be critical for the stable capture and regrowth of MT plus ends against the retrograde flow of lamellar actin that sweeps MTs centripetally. Hence, septins might be part of the mechanisms that enable MTs to reach the front of migrating cells.

In summary, our findings demonstrate that SEPT2/6/7 can directly impact MT dynamics in a manner that depends on their concentration and state of assembly into higher order filamentous structures. Thus, septins could play important roles in the modulation of MT dynamics in health and disease. As septins have been identified in the neurofibrillary tangles and Lewy bodies of Alzheimer’s and Parkinson’s patient brains (Kinoshita *et al.*, 1998; Ihara *et al.*, 2003; Ihara *et al.*, 2007; Shehadeh *et al.*, 2009), filamentous aggregates of septins could have adverse effects on MT dynamics exacerbating the cytotoxicity and pathological progression of neurodegenerative disorders.

## MATERIALS AND METHODS

### Plasmids

The pET-15b plasmid encoding for His-SEPT2-mCherry was constructed and gifted by Dr. Shae Padrick (Drexel University, College of Medicine) and the pnEA-vH and pnCS vectors encoding respectively for His-SEPT2 and SEPT6/7-strep (Mavrakis *et al.*, 2014) have been previously described and provided by Dr. Amy Gladfelter (University of North Carolina, Department of Biology). In the pnCS vector, the N-terminal 19 amino acids (1-57 nucleotides) of human SEPT7 were added using a four-step KAPA Biosystems Site-directed mutagenesis protocol. Briefly, SEPT6/7-strep tagged was amplified with KAPA HiFi HotStart DNA polymerase (KK2502; KAPA BIOSYSTEMS) using the primers 5’-GTAATAATTTTGTTTAACTTTAAGAAGGAGATATACATATGTCGGTCAGTATGG TAGCTCAACAGAAGAA-3’ and 5’-TTCTTCTGTTGAGCTACCATACTGACCGACATATGTATATCTCCTTCTTAAAGTTAAACAAAATTATTAC-3’, to insert the first 12 bp (1-12 bp) at the N-terminus of SEPT7. The amplified PCR product was treated with DpnI (R0176L; New England Biolabs) for 1h at 37°C. DpnI-treated product was heat inactivated at 80°C for 20 min and subsequently was transformed in *E.coli* DH5a cells. The new construct was used as a template and amplified using the primers 5’-GATATACATATGTCGGTCAGTGCGAGATCCG CTGCTATGGTAGCTCAACAGAAGAAC-3’ and 5’-GTTCTTCTGTTGAGCTACCATAGCAGCGGATCTCGCACTGACCGACATATGTATATC-3’, in order to insert the next 15 bp (13-27 bp) of SEPT7. Subsequently, the pcr product was treated as above and used as a template to introduce the next 18 bp (28-45 bp) of SEPT7 using the primers 5’-TGCGAGATCCGCTGCTGCTGAGGAGAGGAGCGTCATGGTAGCTCAACAGA-3’ and 5’-TCTGTTGAGCTACCATGACGCTCCTCTCCTCAGCAGCAGCGGATCTCGCA-3’. Finally, the resulted construct was used as a template and amplified using the primers 5’-TGAGGAGAGGAGCGTCAACAGCAGCACCATGGTAGCTCAACAGA-3’ and 5’-TCTGTTGAGCTACCATGGTGCTGCTGTTGACGCTCCTCTCCTCA-3’, to insert the last 12 bp (46-57 bp) of SEPT7. The pGEX-3x plasmid encoding for GST-tagged EB1 was a gift from Dr. Anna Akhmanova (Utrecht University, Department of Biology). The pET-21a plasmid encoding for recombinant His-tagged EB1-GFP was a gift from Dr. Antonina Roll-Mecak (NIH, Cell Biology and Biophysics) (Vemu *et al.*, 2018). EB3-tdTomato was purchased from Addgene (tdTomato-N1-EB3; Addgene plasmid # 50708; http://n2t.net/addgene:50708; RRID: Addgene_50708) (Merriam *et al.*, 2013). Plasmid encoding rat EGFP-SEPT7 was a kind gift from Dr. Smita Yadav (University of Washington) (Yadav *et al.*, 2017). Plasmid encoding GFP-F-tractin was a kind gift from Dr. Tatyana Svitkina (University of Pennsylvania, Department of Biology) and was cloned into the pEGFP-C1 vector (Johnson and Schell, 2009; Yang and Svitkina, 2019). His-tagged SEPT5 plasmid was constructed by PCR amplifying SEPT5 from pCMV-Myc-tagged SEPT5 (Addgene plasmid # 27272; http://n2t.net/addgene:27272; RRID: Addgene_27272) (Amin *et al.*, 2008) using the primers 5’-ATG GGT CGC GGA TCC GAA ATG AGC ACA GGC CTG-3’ and 5’-CTC GAG TGC GGC CGC ATC ACT GGT CCT GCA TC-3’ and inserting the fragment into the pET28a(+) vector.

### Protein expression and purification

Recombinant SEPT2/6/7 complex was prepared as described before (Mavrakis *et al.*, 2014). The plasmids encoding for His-SEPT2 or His-SEPT2-mCherry and SEPT6/7-strep were co-transformed into *E.coli* BL21 (DE3) (Invitrogen). Bacterial cultures were grown to OD_600_ of 2-3 and induced with 1 mM IPTG for 1h at 37°C (His-SEPT2, SEPT6/7-strep) or to OD_600_ of 0.5 and induced with 0.2 mM IPTG for 16 h at 18°C (His-SEPT2-mCherry, SEPT76/7-strep). Cultures were centrifuged at 4,000 rpm for 20 min at 4°C. Pellets were resuspended in lysis buffer containing 50 mM Tris-HCl pH 8.0, 500 mM KCl, 10 mM imidazole, 5 mM MgCl_2_, 1 mg/ml lysozyme, 1 mM PMSF and 1x bacterial protease arrest cocktail (G-Biosciences; 786-330) and lysed by sonication (10 sets of 15 pulses on ice with 30 s interval between each set). Cell lysates were centrifuged at 13,000 rpm for 30 min at 4°C and passed through a 0.45 µm pore filter. Supernatants were loaded onto gravity flow columns with Ni-NTA agarose beads (745400.25; Macherey-Nagel), which were equilibrated with 10 ml of buffer containing 50 mM Tris-HCl pH 8.0, 500 mM KCl, 10 mM imidazole, 5 mM MgCl_2_, 1 mM PMSF and 1x of the bacterial protease arrest cocktail. Protein complexes were eluted in elution buffer containing 50 mM Tris-HCl pH 8.0, 500 mM KCl, 250 mM imidazole and 5 mM MgCl_2_ and loaded to a StrepTrap HP column (GE Healthcare) equilibrated with buffer containing 50 mM Tris-HCl pH 8, 300 mM KCl and 5 mM MgCl_2_. Protein complexes were eluted in elution buffer containing 50 mM Tris-HCl pH 8.0, 300 mM KCl, 5 mM MgCl_2_ and 2.5 mM d-Desthiobiotin (SIGMA; D1411) and dialyzed overnight at 4°C in BRB80 (80 mM Pipes pH 6.9, 2 mM MgCl_2_, 1 mM EGTA). His-SEPT2-mCherry-SEPT6/7-strep was further purified using an AKTA FPLC system (GE Healthcare) with a Superdex 200 10/300 GL (Amersham Biosciences) gel filtration column.

Recombinant His-tagged EB1-GFP and His-tagged SEPT5 were transformed into *E.coli* BL21 (DE3) (Invitrogen). Bacterial cultures were grown to OD_600_ of 0.6-0.8 and induced with 0.2 mM IPTG for 16 h at 18°C for His-tagged-EB1-GFP or cultures were grown to OD_600_ of 0.4-0.6 and induced with 1 mM IPTG for 5 h at 23°C for His-tagged SEPT5. Cultures were centrifuged at 4,000 rpm for 20 min at 4°C. Pellets were resuspended in lysis buffer containing 50 mM Tris pH 8.0, 150 mM NaCl, 10% glycerol, 1 mM PMSF, 1 mg/ml lysozyme, 10 mM imidazole and 1x Bacterial Protease Arrest protease inhibitor cocktail (G-Biosciences; 786-330) and lysed by sonication (10 sets of 15 pulses on ice with 30 s interval between each set). Cell lysates were centrifuged at 13,000 rpm for 30 min at 4°C and passed through a 0.45 µm pore filter. Supernatants were loaded onto gravity flow column with Ni-NTA agarose beads (745400.25; Macherey-Nagel), which were equilibrated with 10 ml of buffer containing 50 mM Tris pH 8.0, 150 mM NaCl, 10% glycerol, 1 mM PMSF, 10 mM imidazole and 1x Bacterial Protease Arrest protease inhibitor cocktail. Columns were washed with 30 ml washing buffer (50 mM Tris pH 8.0, 300 mM NaCl, 10% glycerol, 10 mM imidazole). Proteins were eluted in elution buffer containing 50 mM Tris pH 8.0, 150 mM NaCl, 10% glycerol and 250 mM imidazole. His-tagged EB1-GFP was dialyzed overnight at 4°C in BRB80 (80 mM Pipes pH 6.9, 2 mM MgCl_2_, 1 mM EGTA) supplemented with 50 mM KCl and was further purified using an AKTA FPLC system (GE Healthcare) with a Superdex 200 10/300 GL (Amersham Biosciences) gel filtration column. His-tagged SEPT5 was dialyzed overnight at 4°C in buffer containing 50 mM Tris pH 8.0, 150 mM NaCl, 10% glycerol.

Recombinant GST and GST-tagged EB1 were transformed into *E.coli* BL21 (DE3) (Invitrogen). Bacterial cultures were grown to OD_600_ of 0.7 and induced with 1 mM IPTG for 5.5 h at 25°C. Cultures were centrifuged at 4,000 rpm for 20 min at 4°C. Cell pellets were resuspended in lysis buffer containing 50 mM Tris pH 8.0, 150 mM NaCl, 2 mM MgCl_2_, 5 mM DTT, 10% glycerol, 0.1% Triton X-100, 1 mM PMSF, 1 mg/ml lysozyme and 1x bacterial protease arrest cocktail (G-Biosciences; 786-330) and lysed by sonication (10 sets of 15 pulses on ice with 30 s interval between each set). Cell lysates were centrifuged at 13,000 rpm for 30 min at 4°C and passed through a 0.45 μm pore filter. Supernatants were loaded onto gravity flow columns with Glutathione agarose beads (16100; Thermo Scientific) equilibrated with 10 ml of buffer containing 50 mM Tris pH 8.0, 150 mM NaCl, 2 mM MgCl_2_, 5 mM DTT, 10% glycerol, 0.1% Triton X-100, 1 mM PMSF and 1x Bacterial Protease Arrest protease inhibitor cocktail. Subsequently, columns were washed with 30 ml washing buffer (50 mM Tris pH 8.0, 150 mM NaCl, 2 mM MgCl_2_, 5 mM DTT, 10% glycerol and 0.1% Triton X-100) and proteins were eluted in elution buffer containing 50 mM Tris pH 8.0, 150 mM NaCl, 2 mM MgCl_2_, 5 mM DTT, 10% glycerol and 10 mM glutathione. All proteins were dialyzed overnight at 4°C in buffer containing 50 mM Tris pH 8.0, 150 mM NaCl and 10% glycerol. Recombinant His-tagged SEPT9_i1 was expressed and purified as described before (Nakos *et al.*, 2019).

### GST-pull down assays

Protein-protein interaction assays were performed between GST or GST-tagged EB1 and recombinant His-tagged SEPT2/6/7 or SEPT5 proteins. GST or GST-EB1 (5 μg) were incubated with 20 μl glutathione agarose beads (16100; Thermo Scientific) for 2 h at 4°C. Subsequently, beads were washed twice with pull down buffer (50 mM Hepes pH 7.4, 150 mM NaCl, 0.1% Triton X-100, 1 mM PMSF, 5 mM DTT, 2 mM EGTA and 10% glycerol) or BRB80 supplemented with 0.1% Triton X-100, 1 mM PMSF and 50 mM KCl. Beads were incubated with His-tagged SEPT2/6/7 or SEPT5 (5 μg) for 2 h at 4°C. Beads were washed five times with pull down buffer or BRB80 (with 0.1% Triton X-100, 1 mM PMSF and 50 mM KCl) before they were resuspended with loading buffer and boiled. Eluted complexes loaded onto 10% SDS-PAGE gels and transferred to a nitrocellulose membrane. Membranes were blocked with 5% non-fat dry milk and 1% BSA for 1 h at room temperature. Membranes were washed with PBS-T (PBS1x/0.1% Tween 20) and incubated with mouse antibody against 6xHis-tag (1:2,000; Qiagen), rabbit antibody against GST-tag (1:10,000; Santa Cruz) or rabbit antibody against SEPT7 (1:10,000; IBL) diluted in PBS-T/2% BSA over night at 4°C. Subsequently, membranes were washed with PBS-T and incubated with anti-mouse or anti–rabbit secondary antibodies (LiCor) for 1 hour at room temperature, before scanning with an imaging system (Odyssey; LICOR).

### Pelleting assay of recombinant SEPT2/6/7

In pelleting assays of recombinant SEPT2/6/7 complexes, increasing concentrations of SEPT2/6/7 (10 nM – 4 μM) were incubated in BRB80 (80 mM Pipes pH 6.9, 2 mM MgCl_2_, 1 mM EGTA) supplemented with 1 mM GTP for 15 min at room temperature. Reactions were placed on top of a cushion buffer and centrifuged at 15,600 x *g* (Optima TL100; Beckman Coulter) for 10 min at 25°C. Equal volumes of supernatant and pellet fractions were loaded onto 8% SDS-PAGE gels which were stained with Coomassie Brilliant Blue. SDS-PAGE gels were scanned and protein bands were quantified with the Odyssey scanning system (LICOR).

### TIRF assays of dynamic MTs

Imaging chambers were prepared and constructed as before (Tanenbaum *et al.*, 2013; Reid *et al.*, 2016; Nakos *et al.*, 2019). TIRF imaging was performed as described before (Nakos *et al.*, 2019). Imaging chambers were prepared by sequential treatment with the following buffers: 1% Pluronic F-127 for 5 min, 5 mg/ml Biotin-BSA (A8549; Sigma-Aldrich) for 5 min, and 0.5 mg/ml Neutravidin (A2666; Invitrogen) for 5 min. GMPCPP-stabilized microtubule seeds (~60-80 nM) were floated into the chamber for 15 min and then washed with blocking buffer (BRB80, 1% Pluronic F-127, 1 mg/ml BSA) for 5 min. GMPCPP MT seeds were prepared by incubating 77% unlabeled tubulin (T240; Cytoskeleton Inc.) with 11.5% biotin-tubulin (T333P; Cytoskeleton Inc.) and 11.5% HiLyte-488-tubulin (TL488M; Cytoskeleton Inc.) or 11.5% HiLyte-647-tubulin (TL670M; Cytoskeleton Inc.). The tubulin mix (20 μM) with 1 mM GMPCPP (NU-405L; Jena Bioscience) in BRB80 (80 mM Pipes pH 6.9, 2 mM MgCl_2_, 1 mM EGTA) was incubated at 37°C for 30 min. After incubation, seeds were diluted in BRB80 and span for 15 min at 100,000 g (Optima TL100; Beckman Coulter). Sedimented MT seeds were resuspended in 60 μl of BRB80. In chambers, MT polymerization was induced by floating a mix of 10 μM tubulin, which contained 6.6% Rhodamine-tubulin (TL590M; Cytoskeleton Inc.) or HiLyte-488-tubulin (TL488M; Cytoskeleton Inc.) in BRB80 with 1 mM GTP, 0.1 mg/ml BSA, 0.1% Pluronic F-127, 0.1% κ-casein, 0.2% methyl cellulose, oxygen scavenging system (0.5 mg/ml glucose oxidase, 0.1 mg/ml catalase, 4.5 mg/ml D-glucose, 70 mM β-mercaptoethanol) and recombinant SEPT2/6/7 or mCherry-SEPT2/6/7.

For TIRF assays with recombinant EB1-GFP and mCherry-SEPT2/6/7, GMPCPP MT bright seeds were prepared as described above by incubating 66.7% unlabeled tubulin (T240; Cytoskeleton Inc.) with 11.5% biotin-tubulin (T333P; Cytoskeleton Inc.) and 21.8% HiLyte-647-tubulin (TL670M; Cytoskeleton Inc.). Microtubule polymerization was induced by introducing a mix of 10 μM tubulin, which contained 4.67% HiLyte-647-tubulin (TL670M; Cytoskeleton Inc.) in BRB80 supplemented with 1 mM GTP, 0.1 mg/ml BSA, 0.1% Pluronic F-127, 0.1% κ-casein, 0.2% methyl cellulose, oxygen scavenging system (0.5 mg/ml glucose oxidase, 0.1 mg/ml catalase, 4.5 mg/ml D-glucose, 70 mM β-mercaptoethanol), 50 mM KCl and recombinant mCherry-SEPT2/6/7 and EB1-GFP proteins. Imaging chambers were sealed with vacuum grease prior to imaging at 37°C. Imaging was performed every 2 s for 15 min.

### Visual septin-tubulin binding assays

Imaging of binding of soluble unpolymerized tubulin to septins by TIRF (Figure S2) was done by adapting a previous MT nucleation assay (Lazarus *et al.*, 2013). Imaging chambers were constructed using acid washed slides and silanized glass cover slips. Acid washed glass coverslips were incubated for 10 min with 20% 3-Aminopropyltriethoxysilane (430941000; ACROS Organics) diluted in acetone and subsequently washed with ddH_2_O for 15 min. Imaging chambers were first incubated with 1% Pluronic F-127 for 5 min. Chambers were washed with BRB80 and then were coated with mouse antibody against 6xHis-tag (1:50; Qiagen) for 10 min. Imaging chambers were washed with BRB80 and blocked with 5% Pluronic F-127 for 5 min. Flow chambers were washed with BRB80-1 mg/ml BSA and subsequently incubated for 15 min with increasing concentrations of recombinant His-tagged SEPT9 or SEPT2/6/7. Unbound proteins were removed from chambers with multiple washes of BRB80-1mg/ml BSA. Subsequently, we floated into the chambers a tubulin mix (3.5 μM), which contained 8.6% Rhodamine-tubulin (TL590M; Cytoskeleton Inc.) in BRB80 supplemented with 2.5 mM GTP, 1 mg/ml BSA, 0.25% Pluronic F-127, 0.1% κ-casein and oxygen scavenging system (0.5 mg/ml glucose oxidase, 0.1 mg/ml catalase, 4.5 mg/ml D-glucose, 70 mM β-mercaptoethanol). Chambers were sealed with vacuum grease and imaging was performed every 2 s for 15 min at 37°C.

### TIRF assays of dynamic MTs with SEPT2/6/7 or actin filaments

GMPCPP-stabilized HiLyte-647-labled MT seeds were made as described above. Imaging chambers were incubated first with 1% Pluronic F-127 for 5 min. Chambers were washed with BRB80 and then were coated with rabbit antibody against SEPT7 (1:10; IBL) for 10 min. Imaging chambers were washed with BRB80 and blocked with 5% Pluronic F-127 for 5 min. Flow chambers were washed with BRB80-1mg/ml BSA and subsequently incubated for 15 min with 800 nM of recombinant mCherry-SEPT2/6/7. Unbound proteins were removed from chambers with multiple washes of BRB80-1mg/ml BSA. Imaging chambers were then incubated with 5 mg/ml Biotin-BSA (A8549; Sigma-Aldrich) for 5 min. Chambers were washed with BRB80-1mg/ml BSA and incubated with 0.5 mg/ml Neutravidin (A2666; Invitrogen) for 5 min. After wash with BRB80-1mg/ml BSA, chambers were incubated with blocking buffer (BRB80, 1% Pluronic F-127, 1 mg/ml BSA) for 5 min. Imaging chambers were washed again with BRB80 containing 1mg/ml BSA and incubated with GMPCPP-stabilized HiLyte-647-labeled seeds diluted in BRB80-1mg/ml BSA for 15 min. Finally, microtubule polymerization was initiated by introducing a mix of 14 μM tubulin, which contained 9% HiLyte-488-tubulin (TL488M; Cytoskeleton Inc.) in BRB80 supplemented with 1 mM GTP, 1 mg/ml BSA, 0.25% Pluronic F-127, 0.1% κ-casein, 0.2% methyl cellulose and oxygen scavenging system (0.5 mg/ml glucose oxidase, 0.1 mg/ml catalase, 4.5 mg/ml D-glucose, 70 mM β-mercaptoethanol). For TIRF assays with EB1-GFP, GMPCPP-stabilized MT seeds and dynamic MT plus ends, were distinguished respectively by brighter and dimmer Hilyte-647 intensity. MT polymerization reaction was additionally supplemented with 75 nM EB1-GFP and 50 mM KCl. Chambers were sealed with vacuum grease and imaging was performed at 37°C, every 2 s for 15 min.

As a control, TIRF assays were performed in the presence of actin filaments. Actin filaments (28 μM) were made by incubating 83% unlabeled G-actin (APHL99; Cytoskeleton Inc.) with 8.5% biotin-G-actin (AB07; Cytoskeleton Inc.) and 8.5% Rhodamine-G-actin (APHR; Cytoskeleton Inc.) in the presence of F-actin buffer (20 mM Hepes pH 7.4, 100 mM KCl, 1 mM MgCl_2_, 0.5 mM ATP and 4 mM DTT) for 1 h at room temperature. Imaging chambers were treated as above but with the following differences: 1) chambers were not coated with an antibody, and 2) 50 nM of Rhodamine F-actin filaments were mixed with GMPCPP MT seeds-HiLyte-647-labeled and incubated for 25 min. MT polymerization reaction was initiated as described above.

### TIRF assays of septin binding to GMPCPP-versus GTPγS-bound MTs

GMPCPP- and GTPγS-stabilized MTs were prepared similar to previous studies (Roth *et al.*, 2018). MTs were made by incubating 77% unlabeled tubulin (T240; Cytoskeleton Inc.) with 11.5% biotin-tubulin (T333P; Cytoskeleton Inc.) and 11.5% HiLyte-647-tubulin (TL670M; Cytoskeleton Inc.) or 11.5% HiLyte-488-tubulin (TL488M; Cytoskeleton Inc.) on ice for 30 min in the presence of 1 mM GMPCPP or GTPγS (371545; EMD Millipore). Subsequently, tubulin mixes were placed at 37°C for 30 min. Then 2 μM of taxol was added to each mix and seeds were further incubated for an additional 5 min at 37°C. MT seeds were diluted in BRB80 containing 2 μM taxol and centrifuged at 100,000 g (Optima TL100; Beckman Coulter) for 15 min. Pelleted MT seeds were resuspended in 60μl of BRB80-2 μM taxol. Imaging chambers were incubated sequentially with 1% Pluronic F-127 for 5 min, 5 mg/ml Biotin-BSA (A8549; Sigma-Aldrich) for 5 min, 0.5 mg/ml Neutravidin (A2666; Invitrogen) for 5 min, ~60 nM of both GMPCPP and GTPγS (mix 1:1) or ~60 nM of GTPγS MT seeds supplemented with 2 μM taxol for 15 min and blocking buffer (BRB80, 1% Pluronic F-127, 1 mg/ml BSA, 2 μM taxol) for 5 min. To test mCherry-SEPT2/6/7 binding to GMPCPP- and GTPγS-stabilized MTs, increasing concentrations of recombinant mCherry-SEPT2/6/7 were incubated for 15 min with GMPCPP and GTPγS MTs in buffer containing BRB80, 0.1 mg/ml BSA, 0.1% Pluronic F-127, 0.1% κ-casein, oxygen scavenging system (0.5 mg/ml glucose oxidase, 0.1 mg/ml catalase, 4.5 mg/ml D-glucose, 70 mM β-mercaptoethanol) and 2 μM taxol. Imaging chambers were sealed with vacuum grease and still images were taken after 15 min of incubation.

For the competition assay, 75 nM of EB1-GFP were incubated for 15 min with GTPγS-stabilized MTs in the presence of increasing concentrations of recombinant mCherry-SEPT2/6/7 in a buffer containing BRB80, 0.1 mg/ml BSA, 0.1% Pluronic F-127, 0.1% κ-casein, 50 mM KCl, oxygen scavenging system (0.5 mg/ml glucose oxidase, 0.1 mg/ml catalase, 4.5 mg/ml D-glucose, 70 mM β-mercaptoethanol) and 2 μM taxol. Imaging chambers were sealed with vacuum grease and still images were taken at 15 min of incubation.

### Primary Cultures and Transfection

Primary rat hippocampal cultures were obtained from the Neuron Culture Service Center (University of Pennsylvania). Hippocampi were dissected from embryonic day 18 pups of mixed genders, obtained from timed pregnant Sprague-Dawley rats. Cells were dissociated by incubation in trypsin-containing media. Cells were plated in Neurobasal medium supplemented with 2% B27 supplement (17504044; ThermoFisher). Cells were cultured in 24 well plates on 12 mm round glass coverslips (Bellco Glass, Inc.), coated with 1 mg/mL poly-L-Lysine (OKK-3056; Peptides International). After 1 day in culture, cytosine d-D-arabinofuranoside (C6645; Sigma) was added at a final concentration of 1.5 μM to inhibit glial proliferation. Cells were maintained at 37°C in a 5% CO_2_ incubator. Neurons were cultured for 1-4 days in vitro (DIV) and transfected with Lipofectamine 3000 (L3000; Invitrogen) for 48 h (DIV 3-6) before live imaging.

### Imaging of rat hippocampal neurons

Live-cell imaging of hippocampal neurons was performed on DIV 3-6, 48 h after transfection. Coverslips were mounted on bottomless 35 mm dishes with a 7 mm microwell using silicone vacuum grease (Beckman Coulter). Conditioned media from the coverslip’s well, along with 2 ml of phenol red-free Neurobasal media supplemented with 2% B27 (17504044, ThermoFisher) and 30 mM HEPES (H0887, Sigma). Dishes were sealed with parafilm. Cells were selected for moderate expression of GFP-SEPT7 or GFP-F-tractin. Neurons were imaged at 30 frames per minute for 3 minutes using total internal reflection fluorescence (TIRF) imaging at 37°C with the TIRF module of the DeltaVision OMX V4 imaging platform.

All TIRF microscopy assays were performed with the DeltaVision OMX V4 inverted microscope (GE Healthcare) equipped with 60x/1.49 NA TIRF objective lens (Olympus), motorized stage, sCMOS pco.edge cameras (PCO), stage top incubator with temperature controller and the softWoRx software.

### Image and statistical analyses

Quantifications of MT dynamics and fluorescence intensities were performed in ImageJ/Fiji. Time-lapse image series were corrected for bleaching using the bleach correction plugin. Kymograph analysis was used to quantify MT dynamics with the KymographBuilder plugin. Growth and depolymerization rates were derived by manual segmentation of the trajectories of growing and shrinking MT plus ends, respectively. The Velocity measurement tool macro was used to calculate rates of polymerization and depolymerization, which excluded the durations of any pausing events. Catastrophe frequencies were measured by dividing the total number of catastrophe events by the total time spent in growing and pausing. Pauses were defined as events without change in the microtubule length for at least 40 s or longer.

Still images at 15 min of incubation were used to quantify the number of tubulin puncta bound to immobilized recombinant SEPT2/6/7 or SEPT9 (Figure S2). Background fluorescence was removed using the subtract background tool (rolling ball radius, 50 pixels) in Fiji. Manual fluorescence thresholding was applied and the analyze particles tool was used to count the number of tubulin puncta.

Quantification of the fluorescence intensity of EB1-GFP (Figure 4B) was performed as previously described (Ramirez-Rios *et al.*, 2016). Background fluorescence was removed with the subtract background tool (rolling ball radius, 20 pixels) and fluorescence intensities of EB1-GFP were measured from kymographs. Segmented lines of a 5-pixel width were drawn manually at the microtubule plus end tips and maximum fluorescence intensities were measured.

Fluorescence intensities of mCherry-SEPT2/6/7 were measured from still images at 15 min (Figure 3D and 4D). Segmented lines of equal width (5 pixels) were drawn manually along the entire length of GMPCPP MT seeds and the entire length of the lattice of plus end segments (Figure 3D) or along the GMPCPP and GTPγS MT seeds (Figure 4D). The average pixel intensity was derived after subtraction of the average pixel intensity of background fluorescence, which was quantified by taking the mean value from three regions that did not contain any MTs. The same method was used to measure the average pixel intensity of EB1-GFP on GTPγS MT seeds (Figure 4F) in the presence of increasing concentrations of mCherry-SEPT2/6/7. Line scans (intensity vs distance plots) in Figure 3C were generated using the plot profile function in Fiji.

Analysis of EB3-tdTomato comet interaction with GFP-SEPT7 or GFP-F-tractin filaments in rat hippocampal neurons was done by making kymographs of neurites from time-lapse TIRF imaging movies using the Segmented Line (15 pixel width) and KymographBuilder tools in Fiji. Kymographs were treated as regions of interest (ROIs), and filaments to track comet pausing or procession were selected from these regions. All comets passing at GFP-F-tractin or GFP-SEPT7 filament were tracked and the number of comets which passed or paused is recorded. A pause in these movies is defined as a halt in comet movement lasting longer than 3 frames (6 seconds). The difference in the rates of EB3-tdTomato movement (ΔVelocity) between before and after pausing at GFP-SEPT7 or GFP-F-tractin were calculated by measuring the rate of movement in the frames immediately before pausing and after resuming growth. For each event of acceleration (V_after_ > V_before_) and deceleration (V_after_ < V_before_), the slower rate was subtracted from the faster and mean ΔVelocity (± SEM) was derived (Figure 5K).

Statistical analysis was performed using GraphPad Prism 6 software. Each data set was tested for normal distribution using the Kolmogorov–Smirnov test. Mean, SEM, SD and *p* values were derived using Student’s *t* test for normally distributed data and the Mann–Whitney *U* test for non-normally distributed data.

## Abbreviations used

MT: microtubules
SEPT2/6/7: septin 2/6/7
SEPT9: Septin 9
SEPT5: Septin 5
EB1: End-Binding protein 1
EB3: End-Binding protein 3
GMPCPP: guanosine-5’-[(α,β)-methyleno]triphosphate
GTPγS: guanosine 5’-O-[gamma-thio]triphosphate
GDP: guanosine diphosphate
GTP: Guanosine-5’-triphosphate
TIRF: total internal reflection fluorescence
DIV: days in vitro

## ACKNOWLEDGMENTS

We thank Drs. Shae Padrick (Drexel University, College of Medicine), Amy Gladfelter (University of North Carolina), Antonina Roll-Mecak (NIH, Cell Biology and Biophysics), Anna Akhmanova (Utrecht University), Smita Yadav (University of Washington) and Tatyana Svitkina (University of Pennsylvania, Department of Biology) for plasmids. All imaging was performed at Drexel University’s Cell Imaging Center. This work was supported by NIH/NIGMS grant GM097664 to E.T.S. The authors have no conflict of interest to declare.

## Supplementary Material

**Figure S1.**
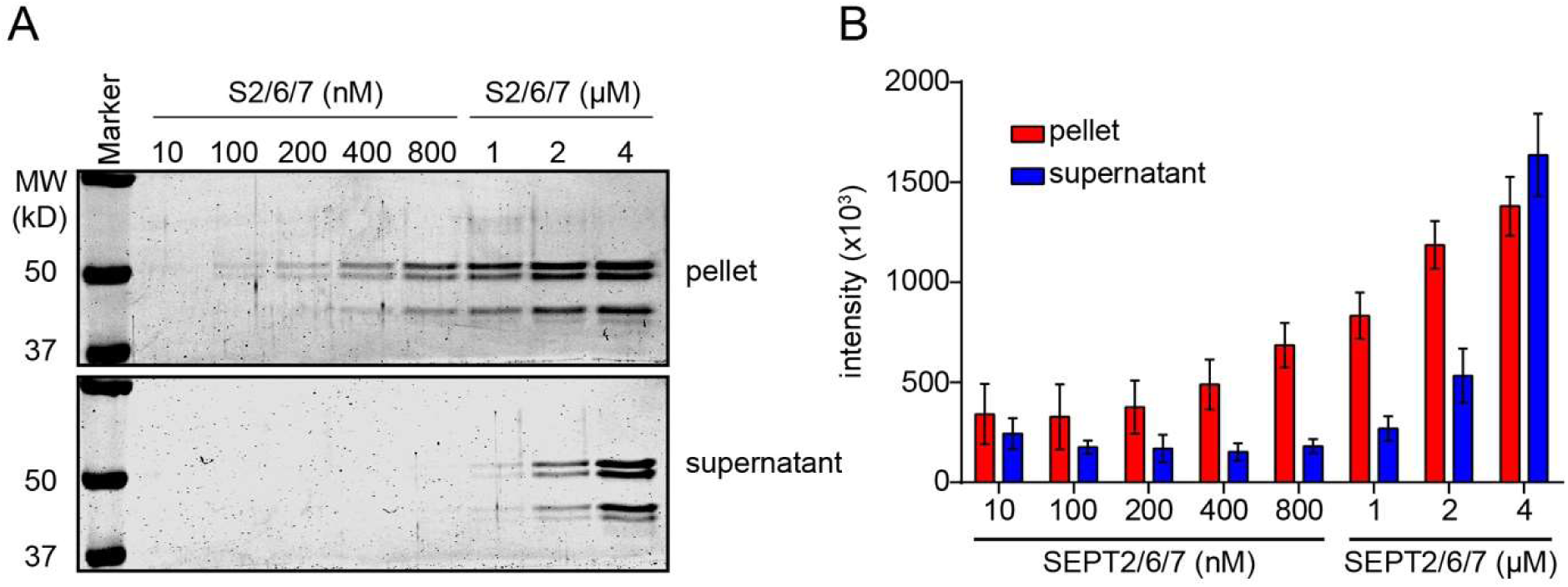
SEPT2/6/7 complexes assemble into higher order filaments in a concentration-dependent manner under conditions of MT polymerization. (A) Coomassie Brilliant Blue-stained SDS-PAGE gels show the pellet and supernatant fractions after sedimentation (15,600 x g) of SEPT2/6/7 at the indicated concentrations. (B) Bar graph shows the mean (± SD) fluorescence intensity of SEPT2/6/7 in pellet (red) and supernatant (blue) fractions at the indicated concentrations.

**Figure S2.**
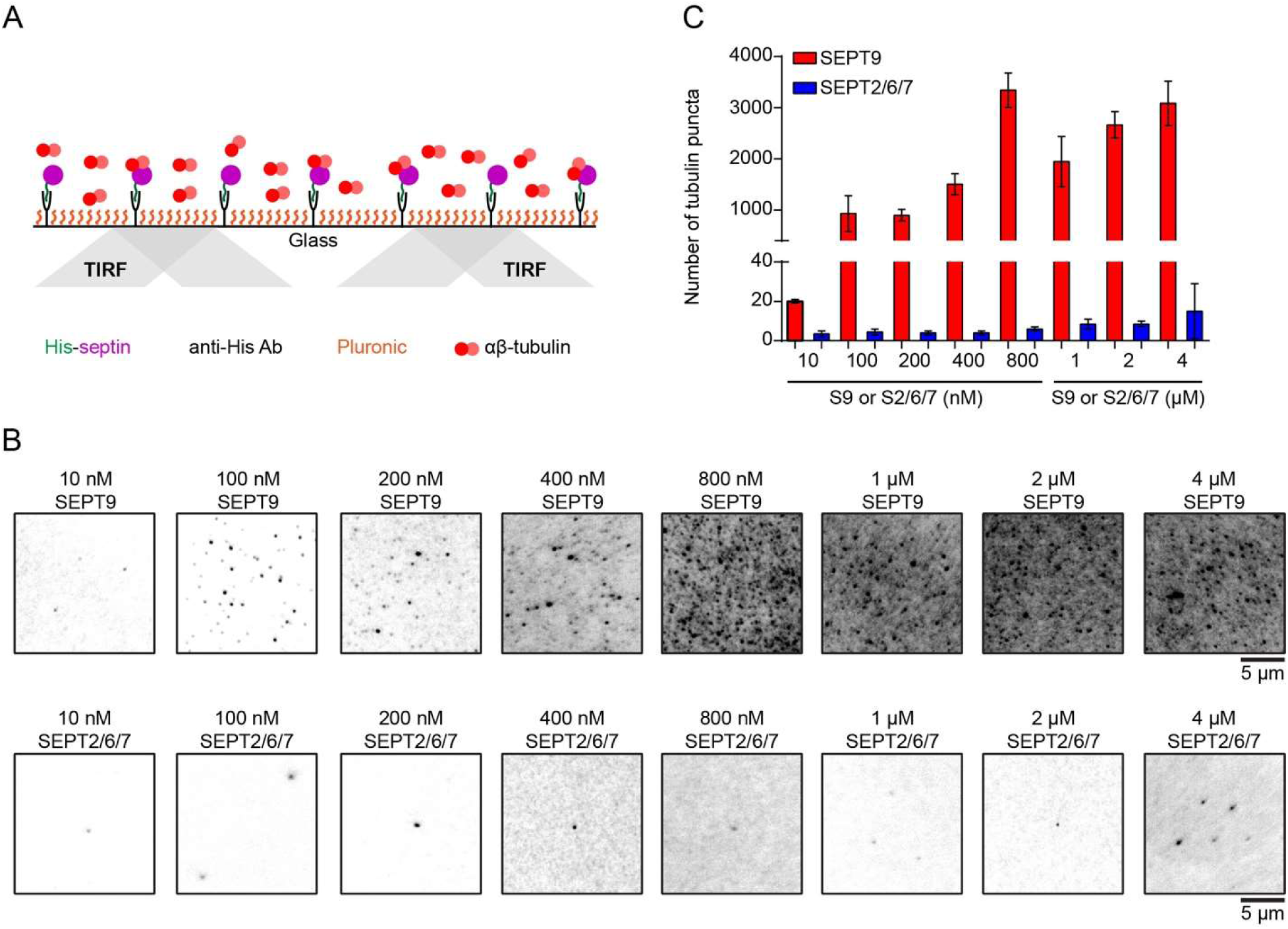
SEPT9 and SEPT2/6/7 differ in their ability to associate with soluble unpolymerized tubulin. (A) Schematic shows the TIRF microscopy assay that used for the MT nucleation assay in the presence of septins. Silanized glass cover slips were used to construct the flow chamber. His-tagged septins were immobilized and the flow chamber was then perfused with Rhodamine-labeled tubulin. (B) Static images of unpolymerized tubulin (inverted monochrome) at 15 min post-tubulin perfusion show that tubulin binding is increased with increasing concentrations of SEPT9 but not with SEPT2/6/7. Note that both SEPT2/6/7 and SEPT9 are not able to nucleate microtubules. (C) Bar graph shows the mean (± SEM) number of tubulin puncta recruited by immobilized SEPT9 (red) and SEPT2/6/7 (blue), 15 min after tubulin perfusion into the chamber.

**Figure S3.**
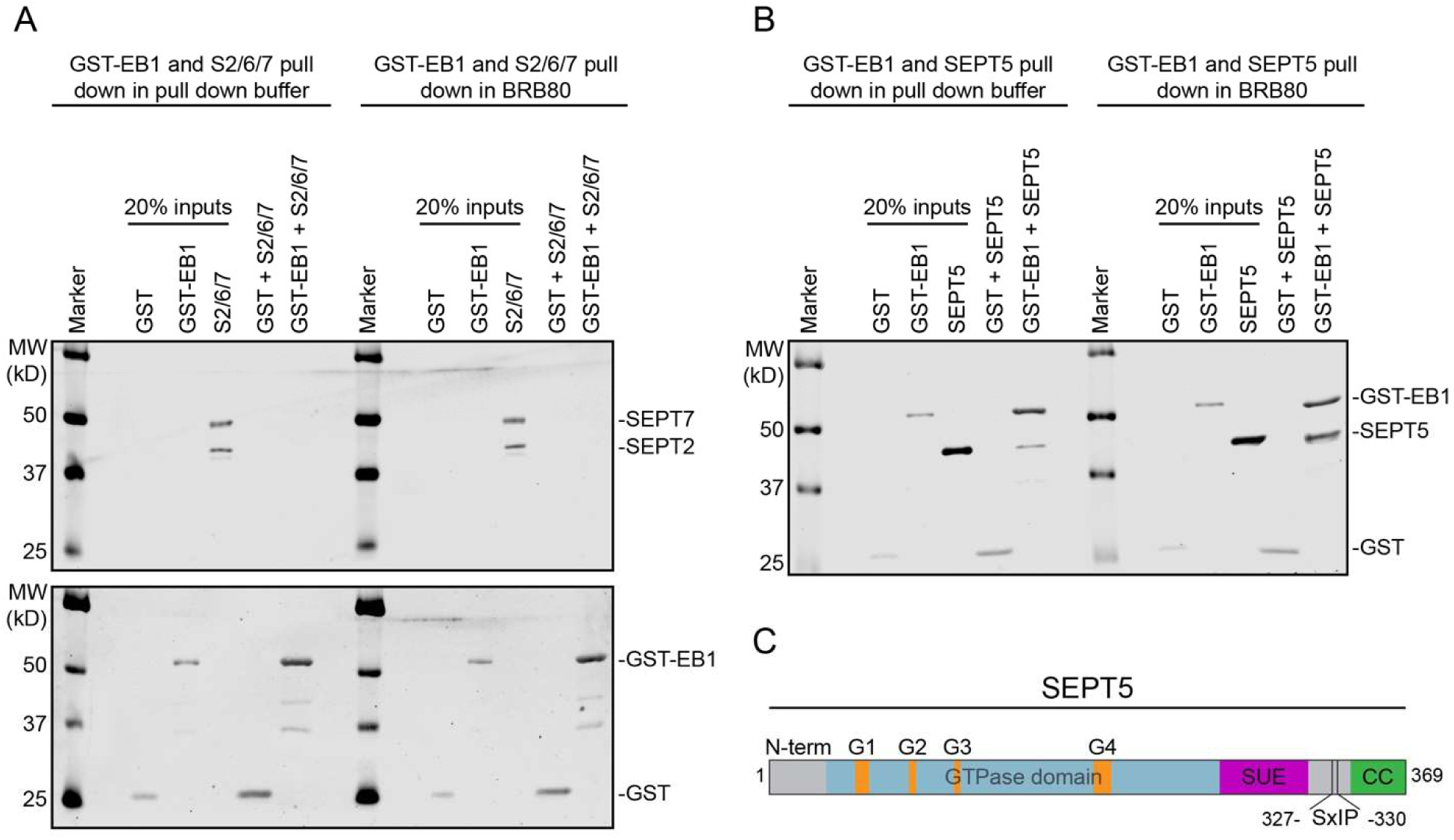
Septin interaction with EB1. (A) Western blots (anti-SEPT7/-His/-GST) of protein binding assays between GST or GST-tagged EB1 and His-tagged SEPT2/6/7-strep. No direct interaction was detected between SEPT2/6/7 and EB1 under two different ionic conditions. (B) Western blots (anti-His/-GST) of protein binding assays between GST or GST-tagged EB1 and His-tagged SEPT5. Note that SEPT5 is physically interact with EB1 under the same ionic conditions that used for SEPT2/6/7. (C) Schematic representation of SEPT5 domains. N-terminus; N-term, G1-G4 motifs; orange, GTPase domain; light blue, Septin Unique Element (SUE); magenta and Coiled Coil (CC); green. SxIP motif locates between the SUE and CC domains (327-330 aa).

## Notes

#### Summary of Updates

Figures and text have been revised and new data have been added.

